# Evaluating genetic ancestry and self-reported ethnicity in the context of carrier screening

**DOI:** 10.1101/208413

**Authors:** Roman Shraga, Sarah Yarnall, Sonya Elango, Arun Manoharan, Sally Ann Rodriguez, Sara Bristow, Neha Kumar, Mohammad Niknazar, David Hoffman, Shahin Ghadir, Rita Vassena, Serena H Chen, Avner Hershlag, Jamie Grifo, Oscar Puig

## Abstract

**Background:** Current professional society guidelines recommend genetic carrier screening be offered on the basis of ethnicity, or when using expanded carrier screening panels, they recommend to compute residual risk based on ethnicity. We investigated the reliability of self-reported ethnicity in 9138 subjects referred to carrier screening. Self-reported ethnicity gathered from test requisition forms and during post-test genetic counseling, and genetic ancestry predicted by a statistical model, were compared for concordance.

**Results:** We identified several discrepancies between the two sources of self-reported ethnicity and genetic ancestry. Only 30.3% of individuals who indicated Mediterranean ancestry during consultation self-reported this on requisition forms. Additionally, the proportion of individuals who reported Southeast Asian but were estimated to have a different genetic ancestry was found to depend on the source of self-report. Finally, individuals who reported Latin American demonstrated a high degree of ancestral admixture. As a result, carrier rates and residual risks provided for patient decision-making are impacted if using self-reported ethnicity.

**Conclusion:** Our analysis highlights the unreliability of ethnicity classification based on patient self-reports. We recommend the routine use of pan-ethnic carrier screening panels in reproductive medicine. Furthermore, the use of an ancestry model would allow better estimation of carrier rates and residual risks.

## BACKGROUND

High throughput technologies like microarrays and next generation sequencing permit the simultaneous interrogation of multiple disease-causing mutations in many genes, allowing expanded carrier screening for almost any disease to be routinely used in medical practice. In contrast, some professional guidelines still recommend carrier screening for certain mutations based on ethnicity[1–3] and physicians commonly use a patient’s self-reported ethnicity to help determine which genetic conditions or mutations to test. When society guidelines recommend expanded carrier screening, they stress that mutation frequencies should be known in the population being tested, so that residual risk in individuals who test negative can be assessed accurately[4,5].

Ethnicity is defined as the membership in a specific group sharing cultural, religious, or racial traits[6], while genetic ancestry refers to the variations in genomic structure among different populations, or the genotypes an individual may have as a result of their ancestors[7]. Because the definition of ethnicity is predicated on shared culture, there is a level of self-identification involved, which may obscure genetic ancestry. For example, an individual may not know about or self-identify with a certain group, despite having an ancestor from it. This issue is further highlighted in admixed populations such as Latinos or African Americans. Different African American groups across the United States have been found to have varying proportions of European contribution to their genome with estimates ranging from 7 to 23% [8]. This genetic heterogeneity means that self-reported ethnicity may not give a complete picture of genetic composition[9].

The incidence of several monogenic diseases varies widely across different populations: Asian Americans account for over 50% of thalassemia cases in the US[10], African Americans are more likely to develop sickle cell anemia[11], and 10-40% of Ashkenazi Jews carry recessive mutations for known Mendelian diseases[12]. As a result, carrier screening programs in the past were selectively targeted to certain racial or ethnic groups[13]. Beyond carrier screening and monogenic diseases, multiple associations between genetic ancestry and clinical outcomes have been observed. For example, lung function in African Americans is inversely correlated with the percentage of African ancestry[14,15], coronary artery calcium and carotid intima media thickness in African Americans and Hispanics vary depending on genetic ancestry and the percentage of European admixture[16], and diabetes among Native Americans is correlated with European admixture[17].

In order to determine the reliability of self-reported ethnicity for clinical decision-making, we compared two sources of self-reported ethnicity, written and verbal, with genetic ancestry as estimated by a statistical model, from a cohort of 9138 patients undergoing genetic carrier screening. Additionally, the relationship between disease carrier status and genetic ancestry within self-reported ethnic groups was studied. Our results indicate that self-reported ethnicity is not a reliable source on which to base clinical decisions.

## MATERIALS & METHODS

### Study Population and Self-Reported Ethnicity

Our study population included 9138 patients undergoing genetic carrier screening, who were referred from fertility specialists, obstetricians/gynecologists, and genetic counselors. Eighty-six percent of the samples were from US clinics, twelve percent from Spanish clinics and the remainder from other countries. Before undergoing in vitro fertilization, couples in fertility clinics were offered CarrierMap (Cooper Genomics, Inc.), a genetic test that determines carrier status for 311 autosomal recessive or X-linked genetic diseases. Informed consent to perform this research was obtained during sample requisition, and confirmed during the genetic counseling session that all patients are offered when CarrierMap results are reported to them. All data presented here is de-identified (HHS 45 CFR part 46.101(b)(4)).

Participants’ ethnicities were self-reported at two separate points in the testing process. The initial report was gathered on the test requisition form, where patients were asked to select all ethnicities that apply from the following list of options (chosen by reviewing the literature): African; East Asian; European; French Canadian; Jewish; Latin American; Mediterranean; Middle Eastern; Native American; South Asian; Southeast Asian; Other (see Supplementary Information for test requisition form). There was also space next to “Other” to write in a response. These responses were mapped to the appropriate category when possible (e.g. Caucasian/White mapped to European). Any participant who selected “Other” without writing in a clarification was excluded from analysis.

The second self-report was made during post-test consultation with a genetic counselor. Standard counseling protocol includes the collection of a complete family history, during which participants were asked to identify their race/ethnicity or the origin of their family. One or more ethnicities from the aforementioned options were then selected based upon the patient’s report. This second source of ethnicity was included in order to determine if there is a difference in what is self-reported depending on the collection technique.

In cases where patients opted out of counseling or were unreachable, a “family history” ethnicity was not generated and the patients were not considered in that part of the analysis. These patients were, however, still included in the comparison between “requisition form” ethnicity and genetic ancestry.

### Genotyping

DNA was extracted from blood or saliva samples, and genomic data was analyzed via Illumina’s Infinium CoreExome-24 v1.0 and v1.1 (catalog ID WG-330-2014 and WG-331-1111, Illumina Inc., San Diego, CA) genotyping platform using standard protocols recommended by the manufacturer.

### Determination of genetic ancestry

In order to estimate the genetic ancestry of our samples, we modified the Expectation Maximization (EM) Algorithm used in FRAPPE[18,19]. We assumed a paradigm where an individual’s genome is divided into segments of different ancestral origin from a set of geographic regions. Our goal was to estimate the proportion of ancestral origin from each geographic region for each individual. Instead of uncovering the ancestral populations from our samples directly, we chose to use an independent source of population allele frequencies to predefine geographic regions at the continental level. It should be noted that this model of genetic ancestry is not intended to fully describe the genetic structure of the human species, as there are significant genetic differences within continental groups. Moreover, there are populations that may be ill described by the geographic groups included in the model. Despite these limitations, the genetic ancestry model is useful as a statistical tool to investigate self-reported ethnic labels in the context of clinical decision-making.

#### Estimation of Allele Frequencies

We obtained allele frequencies for all populations and SNPs in the ALFRED database[20]. This database groups populations by the geographic region where the samples originate. In order to ensure complete allele frequency information, we removed any population that had allele frequency information for fewer than 90% of the total SNPs in the database. Next, we removed populations known to be heterogeneous (African Americans, Hispanics, etc.) and populations that are rare or that could not be easily classified into a single continental group (Yakut, Hezhe). This process left us with a total of 44 populations (Supplementary Table S1). These 44 populations can be further grouped into 8 continental groups: African (AF), Central Asian (CA), East Asian (EA), European (EU), Middle Eastern (ME), Native American (NA), Native Oceanian (NO), and South Asian (SA).

Next, we filtered the list of markers to include only biallelic SNPs existing in Illumina’s Infinium CoreExome-24. We also removed any SNP with a call rate lower than 99.9%. Finally, since the EM algorithm assumes independence of each locus, we used Plink’s pairwise independence command with an R^2^ of .5 and a window size of 1000 to get a final list of SNPs that are in linkage equilibrium[21]. The final set contained 147,550 SNPs.

#### Marker Selection

In order to reduce the full list of SNPs to only those that are informative of ancestry (Ancestry Informative Markers, AIMs), we used Wright’s Fst measure[22]. Instead of considering each population separately, we calculated allele frequencies at the continental level by averaging across populations. Next, we used a two-stage procedure to pick AIMs. First, in order to get a global set of markers, we considered the eight continental groups and picked SNPs that maximize Fst across all groups. Next, in order to ensure a balanced set of AIMs and to distinguish between more closely related groups, we enriched our set with extra SNPs that can differentiate specific groups. We did this by finding SNPs with maximum pairwise Fst, considering each pair of continental groups separately. This process generated a total of 1142 AIMs (Supplementary Tables S2 and S3).

#### Genetic Ancestry Prediction Model

The model was formalized in a similar manner to FRAPPE and ADMIXTURE[18,19]. For a large number of individuals (*I*) and a set of six ancestral populations (*K*), a vector *Q*_*i*_ = (*q*_*i*1_, …, *q*_*ik*_) is estimated for each individual (*i*). Each coordinate in the vector *Qi* is an estimate of the probability that a random allele from *i* originates from population *k*. Our dataset contains genotypes for all *i* at the 1142 AIMs we selected (*M*). Each individual’s genotype is captured in a column vector *g*_*i*_, where each entry *g*_*im*_ holds the number of copies of “allele 1” at marker *m* for individual *i*. The choice of which allele is “allele 1” is arbitrary as long as it is consistent across samples. Departing from FRAPPE, we also have a computed matrix of allele frequencies across populations (*F*). In this matrix, element *f*_*km*_ holds the allele frequency of marker *m* in population *k*.

The log-likelihood of each sample’s vector *Qi* is independent and equal to

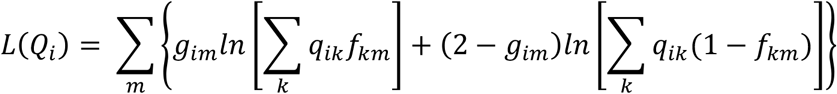

The Expectation Maximization algorithm updates the parameters via the maximization step:

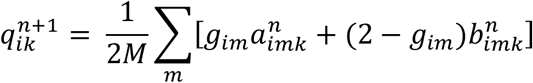

and the expectation step:

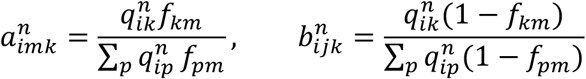

These steps are repeated until the estimation converges. Namely, once:

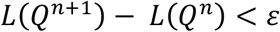

Since we are not estimating the population allele frequencies, we can choose strict stopping criteria (*ε* = 0.01) and still retain an acceptable computation time.

The set of 1142 AIMs was tested on 1000 Genomes Project data by processing genotype data available for 2504 samples (Supplementary Table S4)[23] using the algorithm described above. Genotype data for the 9138 subjects in this study is available upon request or at dbGAP (accession number XXXXX). Scripts used to select the 1142 AIMs and to run the Genetic Ancestry algorithm as well as the Supplementary Data are available at https://github.com/P15/phosphorus-public/tree/master/ancestry.

## RESULTS

The goal of this study was to determine the utility of self-reported ethnicity data for clinical decision-making in the context of genetic carrier screening. In the first step, we selected a set of SNPs that could accurately determine continental genetic ancestry in our patient population. We obtained SNP frequencies from the ALFRED database[24], and through an iterative process, we determined a set of SNPs that could separate the continental groups selected (Supplementary Tables S1 and S2). Supplementary Figure S1 plots the first two principal components of the 44 subpopulations across the 1142 selected SNPs, and shows that six of the eight continental groups are well separated. The Middle Eastern and Central Asian groups are closely related to the European and South Asian groups, respectively, and require an extra set of markers to properly estimate population divergences[25,26] (Supplementary Figure S1). For this reason, we chose not to consider these two groups as separate ancestral populations and removed them from the final estimation.

### Model Validation

In order to validate the genetic ancestry model, we applied it on a set of 2504 samples with known origin from the 1000 Genomes Project[23] (Figure 1 and Supplementary Table S5). The 1000 Genomes Project groups samples by geographic origin and insures that all four grandparents of each sample come from the same group. As expected, the maximum ancestral component of each sample matched the known origin. We conclude that this set of 1142 SNPs correctly estimates continental ancestry in the included populations. These results also validate our approach of using pre-computed population allele frequencies.

**Figure 1.**
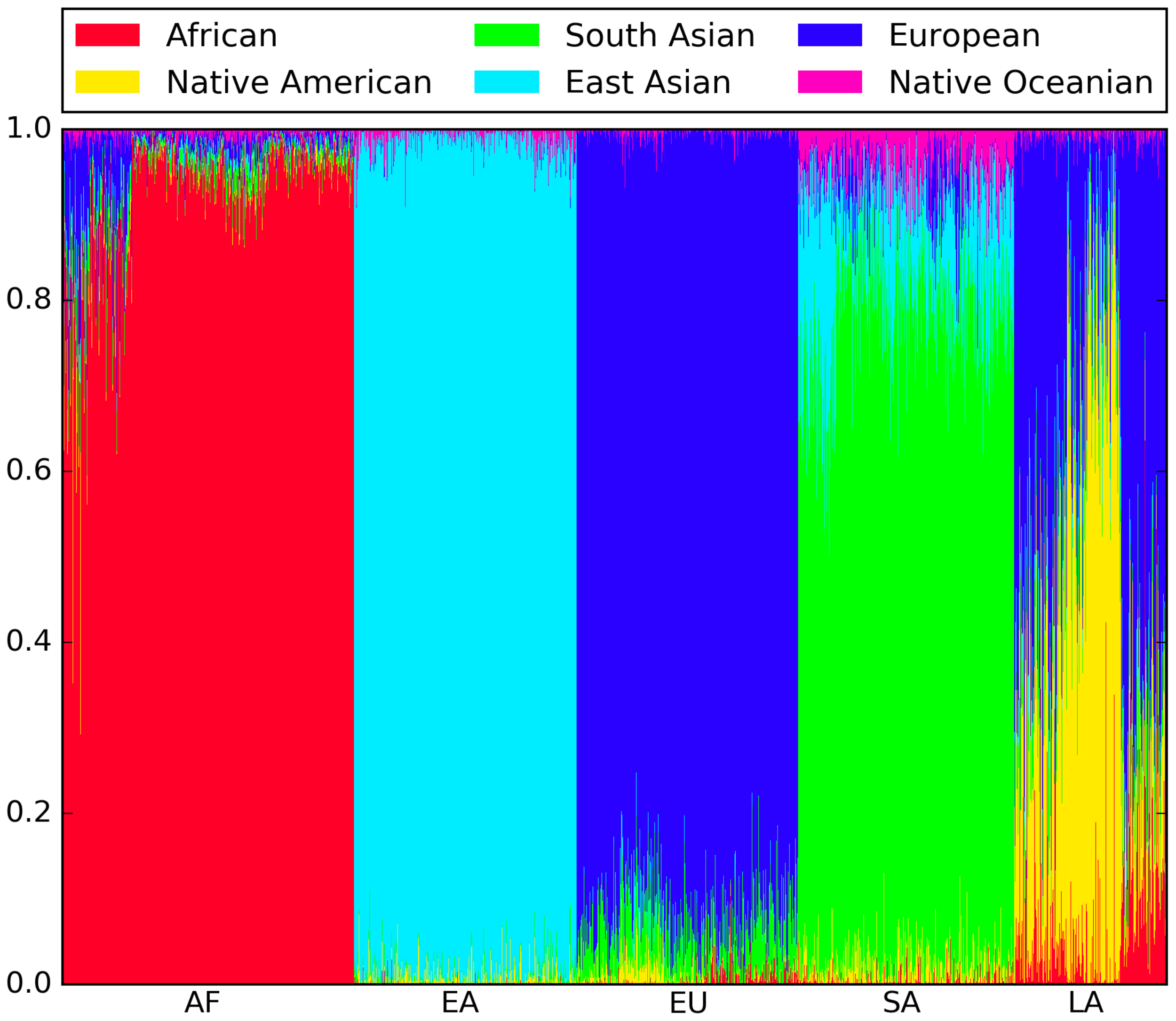
1000 Genomes Project validation results. Each individual is represented as a thin vertical line, where each color shows the proportion of ancestry predicted from each continental group. Individuals are grouped by population: African (AF), East Asian (EA), European (EU), South Asian (SA), and Latin American (LA).

### Ethnicity reported on requisition form vs. family history

We compared self-reported ethnicity from two distinct sources: first, from a requisition form on which patients were asked to check all ethnicities that apply from a provided list and second, from a detailed family history intake performed by a certified genetic counselor. For each ethnic group, we counted the number of patients that selected it on the requisition form, the number of patients that identified it during consults, and the number of patients that did both (Table 1). We excluded all patients that selected ‘Other’ on the requisition form.

**Table 1.**
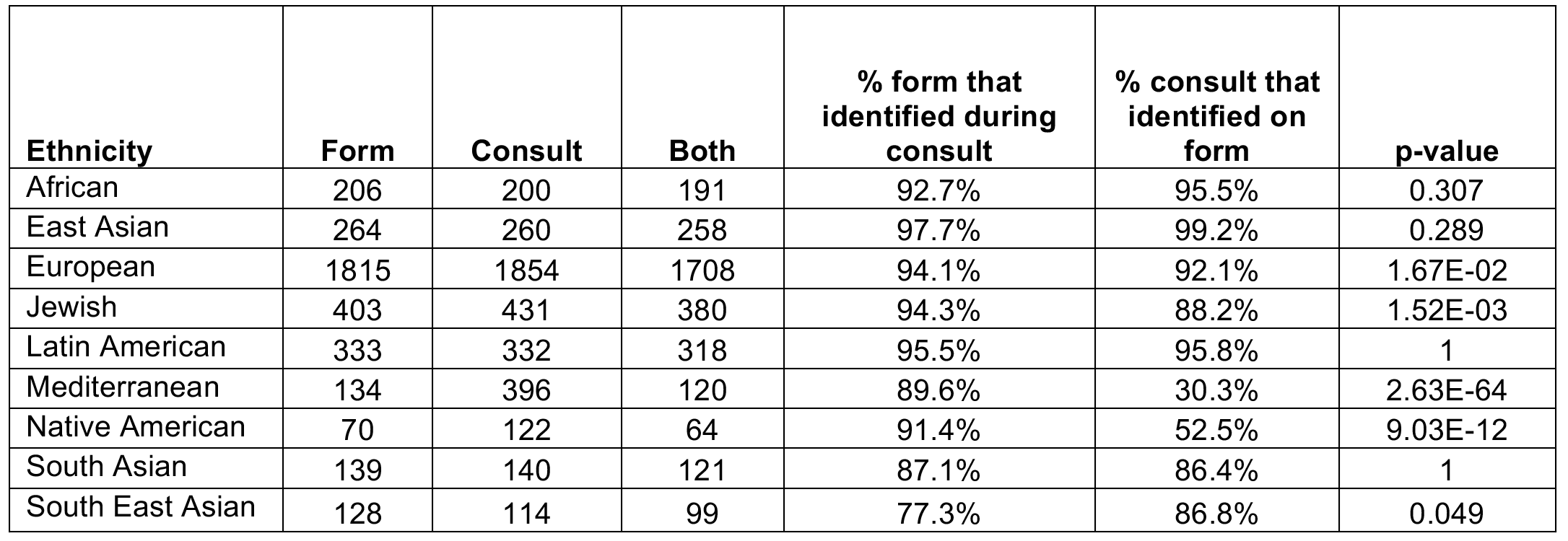
Self-reported ethnicity reported on requisition forms vs. family history discussion. Each row shows counts of the number of patients that selected each ethnicity on the requisition form, reported it during consult, and did both. Additionally the overlap proportions are shown. P-values were computed using McNemar’s test to assess if the proportions are significantly different.

| **Ethnicity** | **Form** | **Consult** | **Both** | **% form that**<br>**identified during**<br>**consult** | **% consult that**<br>**identified on**<br>**form** | **p-value** |
| --- | --- | --- | --- | --- | --- | --- |
| African | 206 | 200 | 191 | 92.7% | 95.5% | 0.307 |
| East Asian | 264 | 260 | 258 | 97.7% | 99.2% | 0.289 |
| European | 1815 | 1854 | 1708 | 94.1% | 92.1% | 1.67E-02 |
| Jewish | 403 | 431 | 380 | 94.3% | 88.2% | 1.52E-03 |
| Latin American | 333 | 332 | 318 | 95.5% | 95.8% | 1 |
| Mediterranean | 134 | 396 | 120 | 89.6% | 30.3% | 2.63E-64 |
| Native American | 70 | 122 | 64 | 91.4% | 52.5% | 9.03E-12 |
| South Asian | 139 | 140 | 121 | 87.1% | 86.4% | 1 |
| South East Asian | 128 | 114 | 99 | 77.3% | 86.8% | 0.049 |

Some of the groups had very consistent patterns of identification across both self-reported sources. For example, 92.7% of patients that selected African on the requisition form identified having African ancestry during the consult, while 95.5% of patients that identified having African ancestry during the consult also selected African on the requisition form. Other groups showed more complex patterns of self-identification. For example, 89.6% of patients that selected Mediterranean on the requisition form also identified it during the consult, while only 30.3% of patients that reported having some Mediterranean ancestry during the consult actually selected it on the requisition form (p-value = 2.63E-64 using McNemar’s test). Similarly, there were significant discrepancies in the requisition form vs. consult for Native American, Jewish and European ethnicities. These differences suggest that the manner in which self-reported ethnicity is collected affects what is reported.

### Self-reported ethnicity vs. genetic ancestry

In order to see the relationship between self-reported and genetic ancestry, we first considered samples that marked only a single ethnicity on the requisition form. We applied our genetic ancestry prediction model to all of these samples and noted the maximum ancestral group (Figure 2A and Supplementary Table S6). We saw a high level of agreement among African and European samples. Of the samples that marked African on the requisition form, 97.2% were predicted to have majority African ancestry by the model. Similarly, of the samples that marked European, 99.3% were predicted to have majority European ancestry by the model. These results are in line with what has been previously reported[27].

The results were quite different when considering Asian continental groups. Of patients that marked only Southeast Asian on the requisition form, 27.5% were predicted to have majority South Asian ancestry by the model, instead of East Asian as expected. At the same time, 8.7% of patients that marked only South Asian on the requisition form were predicted to have majority East Asian ancestry. While this result is consistent with previously published work that shows that the self-reported label “Asian” is concordant with some kind of Asian genetic ancestry, it also suggests that there may be confusion among patients about the distinction between the South, East, and Southeast Asian sub-groups[28].

In order to determine how the source of self-reported ethnicity impacts the genetic ancestry estimate, we repeated this analysis, but this time looked at ethnicity collected during genetic consults. Again, we considered samples that reported only a single ethnicity or country of origin during their consult and noted the maximum ancestral group predicted by the model (Supplementary Table S6). Once again, we saw a high level of agreement between self-reported and predicted ancestry among European and African samples. In contrast to the requisition form, the self-reports from consults among Asian continental groups were mostly concordant with the genetic ancestry estimates. Only 5.6% of patients who self-reported Southeast Asian during the consult were predicted to be have majority South Asian ancestry, with the remaining 93.3% having majority East Asian ancestry. This difference in the genetic ancestry estimates between the two sources of self-reported ethnicity suggests that people may alter what they report depending on the manner in which the report is collected.

### Genetic ancestry prediction in admixed populations

We investigated the predicted genetic ancestry of populations with known admixture (Table 2). Patients that marked only African on the requisition form had on average 79.3% African ancestry and 12.4% European ancestry. Although our samples may include individuals who immigrated from Africa recently, this admixture proportion is in line with previously reported estimates of African Americans[27]. Notably, we found large variability in the predicted proportion of African ancestry in self-identified Africans: from a high of 99% African ancestry to a low of just 1.5% (Figure 2B).

**Table 2.** Mean genetic ancestry component by self-reported ethnicity on requisition form. Each row shows samples based on self-reported ethnicity on requisition forms. Each column shows the average plus minus two standard errors of predicted ancestry proportion across all samples in that self-reported category.

| Requisition Form Ethnicity | African | East Asian | European | Native American | Native Oceanian | South Asian |
| --- | --- | --- | --- | --- | --- | --- |
| African | 79.3%+-1.5% | 1.7%+-0.6% | 12.4%+-1.1% | 1.3%+-0.3% | 1.2%+-0.2% | 4.4%+-0.4% |
| East Asian | 0.1%+-0.1% | 91.7%+-1.7% | 1.2%+-0.5% | 1.2%+-0.2% | 1.2%+-0.2% | 4.9%+-1.4% |
| European | 0.8%+-0.1% | 0.8%+-0.2% | 90.5%+-0.3% | 1.1%+-0.1% | 0.5%+-0.1% | 6.7%+-0.2% |
| Jewish | 3.0%+-0.3% | 1.2%+-0.2% | 79.9%+-0.7% | 1.0%+-0.2% | 1.0%+-0.1% | 14.1%+-0.6% |
| Latin American | 12.0%+-1.0% | 3.3%+-0.4% | 52.1%+-1.2% | 24.4%+-1.2% | 1.0%+-0.1% | 7.5%+-0.3% |
| Mediterranean | 2.6%+-0.5% | 0.9%+-0.3% | 86.4%+-1.0% | 1.4%+-0.4% | 0.7%+-0.1% | 8.2%+-0.6% |
| Native American | 7.3%+-9.2% | 0.7%+-0.5% | 81.4%+-10.2% | 3.8%+-3.5% | 0.6%+-0.4% | 6.5%+-1.8% |
| South Asian | 0.5%+-0.2% | 18.8%+-2.5% | 4.8%+-0.8% | 2.0%+-0.3% | 4.0%+-0.4% | 70.2%+-2.3% |
| South East Asian | 0.5%+-0.3% | 68.3%+-4.3% | 3.0%+-0.8% | 1.7%+-0.3% | 3.4%+-0.4% | 23.4%+-3.9% |
| African & European | 36.4%+-10.2% | 1.1%+-0.6% | 54.8%+-10.6% | 0.8%+-0.4% | 1.0%+-0.5% | 6.1%+-1.5% |
| African & Latin American | 55.0%+-12.6% | 2.4%+-1.1% | 26.4%+-8.0% | 6.3%+-4.9% | 1.5%+-0.8% | 8.7%+-2.6% |
| East Asian & European | 0.4%+-0.3% | 44.3%+-4.8% | 41.8%+-5.1% | 3.8%+-2.0% | 2.1%+-0.7% | 7.9%+-2.5% |
| European & Latin American | 4.2%+-1.3% | 1.8%+-0.5% | 72.7%+-2.6% | 13.4%+-2.4% | 0.8%+-0.3% | 7.4%+-0.9% |

**Figure 2.**
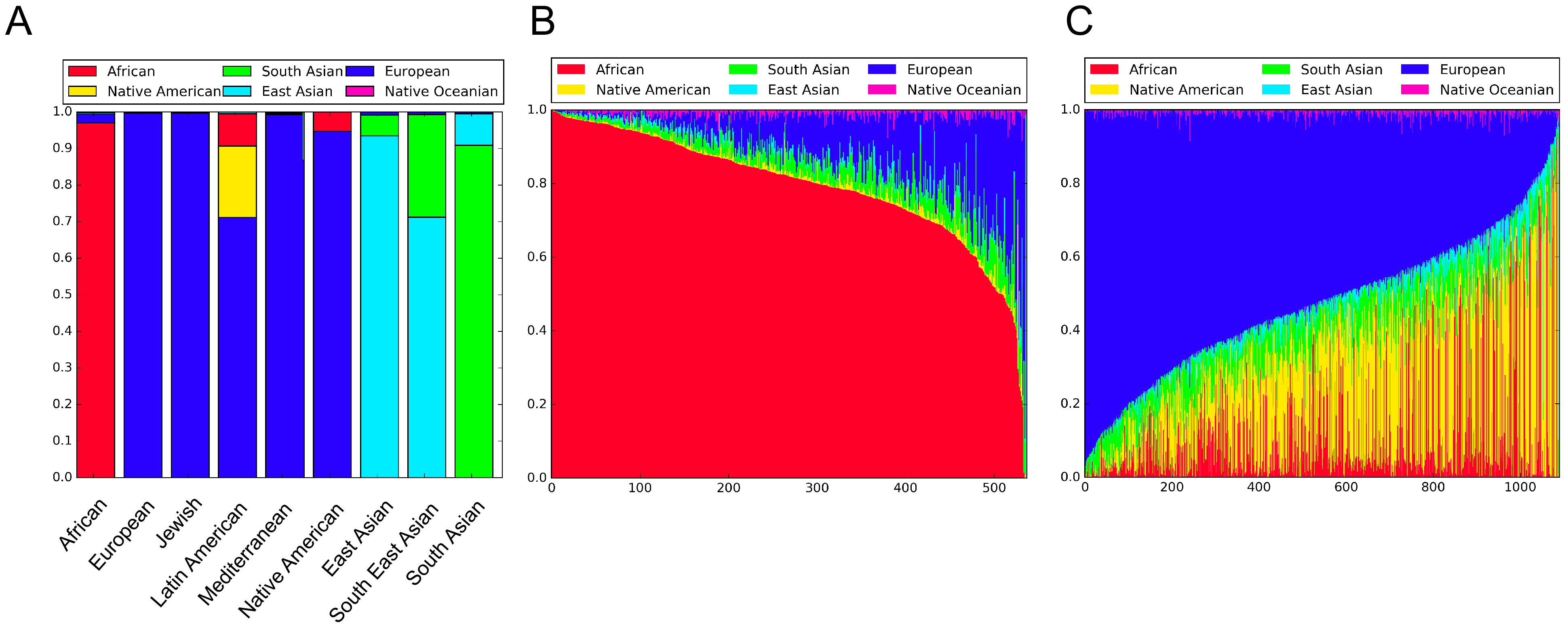
Ancestry model results. (a) Each bar represents all samples within each self-reported ethnicity category. The height of each group shows the proportion of samples that were predicted to have a majority of their ancestry from that group. (b) Individuals who self-reported as only African on requisition forms. Each individual is represented as a thin vertical line, where each color shows the proportion of ancestry predicted from each continental group. (c) Individuals who self-reported as Latin American on requisition forms. Each individual is represented as a thin vertical line, where each color shows the proportion of ancestry predicted from each continental group.

The highest variability was found in patients who marked Latin American on the requisition form. On average, these patients had 52.1% European, 24.4% Native American, and 12.0% African ancestry. These numbers varied widely among individual patients. For example, the Native American component ranged from a high of 90% in some samples to a low of <1% in others (Figure 2C).

We also investigated samples that marked multiple ethnicities on the requisition form and found a wide range of genetic ancestry estimates (Table 2). For example, on average, patients that marked both European and East Asian on the form were predicted to have 41.8% European and 44.3% East Asian ancestry. Among the individual samples however, the East Asian component ranged from a low of 8.8% to a high of 58%. Finally, we considered samples with at least 20% predicted ancestry from both the European and East Asian continental groups (88 subjects). Of these samples, 29 (33%) reported both European and Asian on the requisition form, while 15 (17%) reported only Asian, 12 (13.6%) reported only European, and 32 (36.4%) reported something else (Supplementary Table S7).

### Carrier risk in admixed populations

In order to better understand how genetic ancestry estimates within self-reported ethnic groups impact clinical estimates of risk, we investigated the relationship between carrier status of cystic fibrosis, sickle cell anemia, and GJB2-related nonsyndromic hearing loss and predicted ancestry among self-reported African and Latin American patients. These diseases were chosen because they are common and strongly associated with specific ancestral groups. We found significant differences in carrier risk based on genetic ancestry (Supplementary Table S8). For example, the mean African ancestry of Latin American patients identified as carriers of sickle cell anemia is 30.3%, while the mean African ancestry of non-carriers is 11.5% (P<0.0001, Mann-Whitney’s U test). Similarly, the mean European ancestry of Latin American patients identified as carriers of cystic fibrosis is 59.4% while the mean European ancestry of non-carriers is 51.9% (P=0.03). Finally, the mean European ancestry of African carriers of GJB2-related nonsyndromic hearing loss is 23.0%, while the mean European ancestry of non-carriers is 12.2% (P =0.01).

In order to see how carrier rates differ based on genetic ancestry, we computed the 80^th^ percentile of European and African ancestry proportion among self-reported Latin Americans and Africans. We then computed carrier rates among individuals above and below this 80^th^ percentile threshold and found significant differences in carrier rates of these diseases (Table 3).

**Table 3.** Carrier rates by genetic ancestry. Differences in carrier rates of sickle sell anemia, cystic fibrosis, and GJB2-related nonsyndromic hearing loss by proportion of African and European ancestry among self-reported Africans and Latin Americans. For each group, an ancestry threshold was chosen by computing the 80^th^ percentile of ancestry proportion. For example the 80^th^ percentile of African ancestry among Latin Americans is 18.68%. Thus, the carrier rate of sickle cell anemia of Latin Americans below this threshold is 1.261% and the carrier rate above it is 4.587%. P-values were computed using Fisher’s exact test.

| **Sickle Cell Anemia** |  |  |  |  |
| --- | --- | --- | --- | --- |
|  | **carrier rate <**<br>**threshold** | **carrier rate >**<br>**threshold** | **p-value** | **African Ancestry Threshold**<br>**(80th percentile)** |
| African | 6.760% | 15.741% | 0.00605 | 93.439% |
| Latin American | 1.261% | 4.587% | 0.00372 | 18.677% |
| **Cystic Fibrosis** |  |  |  |  |
|  | **carrier rate <**<br>**threshold** | **carrier rate >**<br>**threshold** | **p-value** | **European Ancestry Threshold**<br>**(80th percentile)** |
| African | 1.631% | 6.482% | 0.01095 | 19.928% |
| Latin American | 1.606% | 3.670% | 0.06128 | 68.715% |
| **GJB2-Related Nonsyndromic Hearing Loss** |  |  |  |  |
|  | **carrier rate <**<br>**threshold** | **carrier rate >**<br>**threshold** | **p-value** | **European Ancestry Threshold**<br>**(80th percentile)** |
| African | 1.399% | 5.556% | 0.01870 | 19.928% |
| Latin American | 3.440% | 3.670% | 0.83750 | 68.715% |

For example, the carrier rate of cystic fibrosis among Latin Americans with less than 68.7% European ancestry is 1.6%, while the carrier rate of those with more than 68.7% European ancestry is 3.7%. To highlight the significance of this difference, if we assume a detection rate of 72% (based on the ACMG panel of 23 common CFTR mutations), in the case of a negative screen, the residual risk of actually being a carrier is 1/220 in Latin Americans with less than 68% European ancestry and 1/94 in Latin Americans with a higher European ancestry proportion. A similar situation is observed for Africans with sickle cell anemia, where the carrier rate varies from 6.7% to 15.7%, depending on whether they have less or more than 93% African ancestry. Finally, the carrier rate of GJB2-related nonsyndromic hearing loss among Africans varies from 1.4% to 5.6%, depending on whether they have less or more than 20% European Ancestry.

## DISCUSSION

We compared two sources of self-reported ethnicity to genetic ancestry as predicted by a validated statistical model. First we measured concordance between a written self-report on a requisition form and a verbal report during a genetic counseling session. While for most ethnicities there was concordance between them, for ethnicities like Mediterranean, Native American, and Southeast Asian, responses on the requisition form and during genetic counseling were different. We also observed differences between self-reported ethnicity on the requisition form and genetic ancestry in South Asians and Southeast Asians. Notably, these differences were mitigated when looking at self-reported ethnicity during genetic counseling consults. The discrepancies imply that there is confusion about the meaning of the different labels, impacting our ability to rely on self-reported ethnicity, which is in line with prior reports[29,30].

In the realm of carrier screening, the main consequence of inaccurate classification is miscalculation of reproductive risk. Our results show that carrier rates and residual risks are dependent on genetic ancestry in admixed populations (Table 3). For example, for Latin Americans, the carrier rate of cystic fibrosis varies from 1.6% to 3.7% depending on the percentage of European ancestry, and the carrier rate of sickle cell anemia varies from 1.3% to 4.6% depending on the percentage of African ancestry. Additionally, we found that in admixed populations, the proportion of genetic ancestry estimated from each continental group as well as what ethnicities are self-reported, varies by individual. As such, it is erroneous to assume that genetic disease risks affect admixed populations in a uniform manner.

Carrier rates and residual risk estimates vary significantly depending on which source is used to account for population differences. Self-identification does not always paint a complete picture, as there may be uncertainty about family origins, confusion about labels, and identification or lack thereof with a particular group due to personal or cultural reasons. Since mutation allele frequencies are linked to genetic ancestry, using self-identified ethnicity to select panel content or to adjust for risk will lead to errors, which may impact clinical decisions. Despite recommendations from professional societies to base some carrier screening decisions on ethnicity, this study suggests that in order to ensure that carriers of severe genetic disorders are not missed, expanded pan-ethnic carrier screening panels should be utilized. Pan-ethnic panels are able to detect mutations regardless of minor allele frequency (low or high), and include all mutations described for a given disease, present in different populations. Additionally, targeted next generation sequencing can be used to detect novel or poorly described mutations in genes of interest. While, pan-ethnic panels have some disadvantages[31,32], given the unreliability of self-reported ethnicity and the goal of providing couples with information to optimize pregnancy outcomes, pan-ethnic panels represent the most comprehensive approach.

A second conclusion from this study is that, since carrier rates, detection rates and residual risks vary based on ancestry, in order to accurately counsel patients about their reproductive risks, genetic ancestry should be determined in clinical practice. Prior reports also describe sets of Ancestry Informative Markers (AIMs) that could be used to determine genetic ancestry, although further work is needed to ensure that allele frequency estimates are representative of all populations and that the ancestry estimates are meaningful for all individuals[33–38].

Limitations of the present study include that the data was studied retrospectively using the records of a clinical carrier screening company. Self-reported ethnicity from requisition forms was entered into the database during sample accessioning on a rolling basis. It is possible that some of the records were entered incorrectly or later modified by genetic counselors. To ensure accurate measurement, we manually reviewed several hundred scanned requisition forms: both randomly and in cases where discordance between self-reports and genetic ancestry was found. Reviewing records randomly showed that less than 1% of the self-reported labels were incorrect in the database. Another limitation is that our genetic ancestry model is based on allele frequency estimates from a limited sample size and assumes that grouping individuals by continent provides meaningful estimates of origin. A better way would be to account for subtle differences in genetic ancestry among local subpopulations, however, this would require expanded screening platforms and increased resources. Despite these limitations, we were able to correctly estimate the origin of validation samples from the 1000 Genomes Project and we found a strong relationship between carrier rates and ancestry proportion in admixed populations. This shows that the ancestry model provides meaningful information. Further work with larger cohorts is needed to refine the ancestry model and to measure the relationship between carrier rates and genetic ancestry for more diseases. Additionally, in diseases where environmental factors are known to play a part, further work is needed to understand how much impact ancestry has versus social and cultural influences of an individual’s self-identified group.

## CONCLUSIONS

Estimates of carrier rates and residual risks depend on whether self-reported ethnicity or genetic ancestry is used to account for population differences. Furthermore, self-reports are not reliable for clinical decision-making. In order to mitigate the risk of ordering the wrong testing panel, we recommend the use of expanded pan-ethnic carrier screening panels. Additionally, in order to accurately estimate carrier rates and residual risks, we recommend the use of a genetic ancestry model in clinical genetic testing.

## DECLARATIONS

### Ethics approval and consent to participate

Institutional Review Board to consent for this research was obtained from ASPIRE (IRB-R-004). All data presented here is de-identified (HHS 45 CFR part 46.101(b)(4)).

### Availability of data

Genotype data for the 9457 subjects in this study is available upon request or at dbGAP (accession number XXXXX). Scripts used to select the 1142 AIMs and to run the Genetic Ancestry algorithm as well as Supplementary Data are available at https://github.com/P15/phosphorus-public/tree/master/ancestry.

### Authors’ contributions

Roman Shraga, Arun Manoharan and Oscar Puig participated in study design and data analysis. Roman Shraga, Sarah Yarnall, Sonya Elango, Arun Manoharan, Sally Ann Rodriguez, Sara Bristow, Neha Kumar, Mohammad Niknazar, David Hoffman, Shahin Ghadir, Rita Vassena, Serena H Chen, Avner Hershlag, Jamie Grifo and Oscar Puig participated in data collection, interpretation of results and drafting the manuscript.

## Acknowledgements

We thank Ed O’Neill for editorial comments in the manuscript.

## Supplementary Figure S1 legend

Plot of the first and second principal components obtained by Principal Component Analysis on 44 geographic groups (described in Supplementary Table S1) and 1142 AIMs. Each geographic group is shown as a point and is colored according to the continental group to which it belongs. The plot illustrates that the AIMs separate most continental groups well, but the Middle Eastern and Central Asian groups do not form distinct clusters.

